# Dopamine Compensates for Amyloid-Induced Default Mode Network Dysfunction to Support Learning

**DOI:** 10.64898/2026.07.07.736872

**Authors:** Joseph Giorgio, Thomas M. Morin, Hsiang-Yu Chen, Anne S. Berry, Michael Breakspear, William J. Jagust

## Abstract

Throughout the preclinical phase of Alzheimer’s disease (AD) β-amyloid (Aβ) accumulates preferentially within the default mode network (DMN), yet the functional and behavioural consequences of this pathological burden remain poorly understood. Using task-based fMRI combined with Aβ, tau, and dopamine PET in cognitively normal older adults, we show that Aβ burden impairs learning independent of tau, but this learning performance is recovered with higher dorsolateral striatal dopamine synthesis capacity. Investigating the neural mechanisms that support this learning, we observe that Aβ positive individuals show attenuated DMN activity to error related feedback, a metric that relates to poorer learning. When estimating the effective connectivity during feedback, computational modelling reveals that Aβ induces dis-inhibition of the DMN during error processing. Critically, dopamine synthesis capacity in the dorsolateral striatum rebalances effective connectivity between the DMN and frontostriatal network, thereby opposing Aβ related disruption. These findings establish a systems-level framework in which Aβ impairs learning by disrupting dynamic DMN modulation during feedback, a disruption for which dopaminergic function can partially compensate. This suggests that learning in the presence of Aβ may be subserved by dopamine-dependent network rebalancing, a candidate mechanism of cognitive resilience to support learning in preclinical AD.

## Introduction

A fundamental function of the human brain is to build models of our environment and compare new sensory stimuli against these existing models ^1^. When faced with new sensory evidence that contradicts our existing model, prediction errors shift our previous beliefs, which in turn update internal models. This mechanism defines how the brain adapts when learning and is regulated by neuromodulatory systems centred on dopamine^2,3^. How this critical process that supports effective learning is impacted by Alzheimer’s disease (AD) is not known^4^.

Late-onset sporadic AD is characterised by a stereotyped progression of the accumulation of pathological forms of tau in medial temporal lobe, the deposition of β-amyloid (Aβ) in association cortex, and the subsequent aggregation of tau throughout the association cortex which coincides with the transition to clinical phases of AD ^5,6^. This symptomatic phase is preceded by a protracted preclinical phase where Aβ gradually accumulates within regions overlapping with the default mode network (DMN) in the absence of overt cognitive deficits ^7–9^. Recent evidence suggests that individuals in the preclinical phase of AD undergo subtle cognitive changes. While episodic memory declines substantially in the presence of Aβ and neocortical tau^10^, Aβ deposition in preclinical AD prior to advanced tau burden is associated with disturbances in executive functioning ^11–13^. Longitudinally, preclinical Aβ deposition attenuates learning effects over time, with a reduction of expected increases in performance during repeated cognitive testing ^14,15^. These findings suggest in the preclinical AD phase cognitive processes more closely related to executive functioning and learning may be more sensitive to early Aβ than hippocampal-dependent episodic memory.

Some individuals burdened with AD pathology maintain higher-than-expected performance (i.e. cognitive resilience)^16^ . While the cortical processes underlying this cognitive resilience remain poorly characterised, the compensatory action of neuromodulatory systems such as the cholinergic, noradrenergic, and dopaminergic pathways have been implicated ^17^. However, the role of striatal dopamine synthesis capacity in maintaining better cognitive performance is unclear. Prior work in older adults has suggested a positive role for higher striatal dopamine synthesis capacity on working memory.^18,19^ , whereas other work has suggested a negative role in relation to cognitive flexibility^20^. How dopamine synthesis capacity specifically relates to learning and prediction error processing in older adults with AD pathology has not been established.

Abundant evidence demonstrates prediction-error activity in frontostriatal circuits comprising the ventral striatum and ventromedial prefrontal cortex, implicating this circuit in feedback learning, including predictive error processing ^21–23^. Beyond frontostriatal brain function, the DMN also contributes to learning through dynamic responses to prediction error by comparing internally generated probabilistic models against incoming information ^24,25^. Notably, prior work implicated connectivity between the DMN and the dopaminergic midbrain during surprising events; suggesting that model updating in response to prediction errors relies on dynamic coupling between the DMN and striatal regions of the dopaminergic reward system^25^. How these networks activate and co-activate to support learning under the pathological influence of Aβ in preclinical AD is yet to be fully established.

Here, we investigated the neural computations that support learning via feedback and how this relates to AD pathologies and dopamine synthesis capacity. Using task-based fMRI, younger adults and cognitively normal older adults with varying degrees of AD pathology were imaged while performing a probabilistic reward learning task paired with an encoding phase to probe episodic memory. This paradigm engages two cognitively separable processes, reinforcement learning and episodic memory. Prior work has characterised how AD pathology and dopamine synthesis capacity interact to disrupt subsequent memory for this task^26^. Here we focus on the complementary learning process, examining how prediction error signals, agnostic to outcome valence, modulate neural activity during feedback and how Aβ burden and dopamine synthesis capacity jointly shape this modulation. Based on the topography of Aβ and the role of frontostriatal networks in feedback learning we aimed to assess how the DMN and frontostriatal networks interact when processing feedback and the role of Aβ and dopamine in these interactions. As prior literature implicates dorsolateral striatum (DLS) dopamine in stimulus-response associations ^27,28^ we focus our analysis on dopamine synthesis capacity in this region. Using system level analysis, we hypothesise that Aβ pathology disrupts the normative DMN processing of error feedback, which in turn impairs learning. Furthermore, we anticipate that DLS dopamine acting through frontostriatal effective connectivity with the DMN may counteract this Aβ related disruption to support effective learning in preclinical AD.

## Results

### Study Overview

Fifty cognitively normal older adults (65–85 years) and thirty healthy younger adults (18–35 years) participated in the study. All participants provided written informed consent in accordance with procedures approved by the Institutional Review Board at the University of California, Berkeley. Older participants were recruited through the Berkeley Aging Cohort Study. During fMRI scanning, participants performed a probabilistic reward-learning task for which methods have been published previously (see Morin 2026^26^, **Figure 1**). Briefly, participants were instructed (**Supplementary Methods**) that they owned a portfolio of houses and were working with three real estate agents to sell them. The agents differed in their typical performance with one agent tending to sell houses for a profit (“reward agent”), one consistently breaking even (“neutral agent”), and one tending to sell houses at a loss (“loss agent”). During the cue phase, participants indicated the identity of the agent by pressing one of three buttons. This was followed by presentation of the house being sold (target phase) and outcome feedback indicating the result of the sale (gain, break-even, loss). Outcomes were predetermined according to fixed probabilities associated with each agent (reward agent: 80% profit, 20% loss; loss agent: 20% profit, 80% loss; neutral agent: 100% break-even) and were independent of participant responses. Participants gained or lost real money based on the outcome of each sale, as results were deterministic all participants received $30 if they responded to each trial.

**Figure 1.**
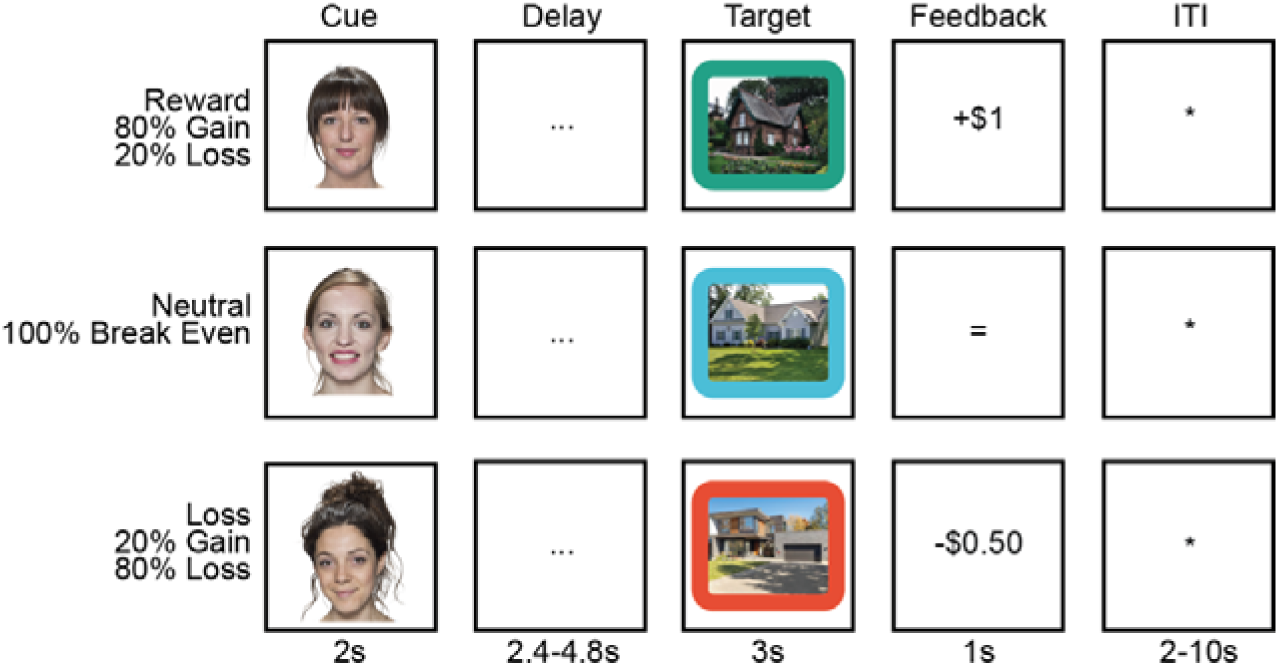
Real Estate Selling Task. During fMRI scanning, subjects completed a Real Estate Selling Task. The initial cue presented the face of a realtor, participants then indicated if they predicted the realtor would sell a house at a gain, loss, or break even. Participants were then presented a house and feedback indicating if the realtor sold the house at a gain, loss, or broke even. Face stimuli taken from DeBruine, L. & Jones, B. (2017). Face Research Lab London Set. figshare. doi:10.6084/m9.figshare.5047666, licensed CC-BY 4.0.

As part of the longitudinal Berkeley Aging Cohort Study protocol, older adults underwent extensive neuropsychological assessments which included the digit span task to assess working memory, and the California verbal learning task delayed forced recall to assess episodic memory. In addition, a subset of participants also underwent PET imaging to quantify molecular markers relevant to learning and neurodegeneration. In older adults dopamine synthesis capacity was measured using [^18^F]fluoro-L-m-tyrosine (FMT) PET (n = 41), tau pathology was assessed using [^18^F]flortaucipir (FTP) PET (n = 41), and amyloid-β (Aβ) burden was measured using [^11^C]Pittsburgh Compound-B (PiB) PET (n = 45). As the aim of this study was to characterise the role of AD pathology and dopamine in learning stimulus response associations, we extracted three measures of interest from the PET imaging. From the FMT-PET we extracted signal of dopamine synthesis capacity from the dorsolateral striatum (DLS), from the FTP-PET we extracted standardised uptake volume ratio (SUVR) from the temporal meta region of interest ^29^, and we assigned Aβ positivity as a PiB distribution volume ratio >1.065^30^.

Of the total sample, 44 older adults had in-scanner behavioural performance that could be estimated (excluded participants responded with fingers placed incorrectly on button box). Of these, 33 had all three PET imaging variables available (39 with FMT and PiB; 44 with PiB). 42 of these older adults had fMRI data that were successfully pre-processed using fMRIPrep^31^ and from this sample all had PiB PET available and 37 had PiB and FMT available. All 30 younger adults had behavioural and fMRI data available (**Supplementary Table 1**).

### Learning Performance

We derived subject-level learning curves by computing the trial wise Kullback–Leibler divergence between participants’ responses to each face stimulus and the associated feedback distribution (see Methods). As the stimulus–response mapping of the neutral realtor was deterministic, these learning curves were calculated only for the winning and losing realtors. To derive a subject-specific metric of learning we took the average of the integral of the winning and losing realtor learning curves. This learning measure approaches 0 when the response distribution exactly matches the feedback distribution and is larger when the distributions diverge (i.e. larger values represent poorer learning) (see **Supplementary Figure 1** for exemplar learning curves and response distributions). Comparing learning performance between age groups revealed a significant difference, with younger adults demonstrating superior learning relative to older adults (t(72) = −3.2, p = 0.002, Cohen’s d = −0.81) (**Figure 2a.).**

**Figure 2.**
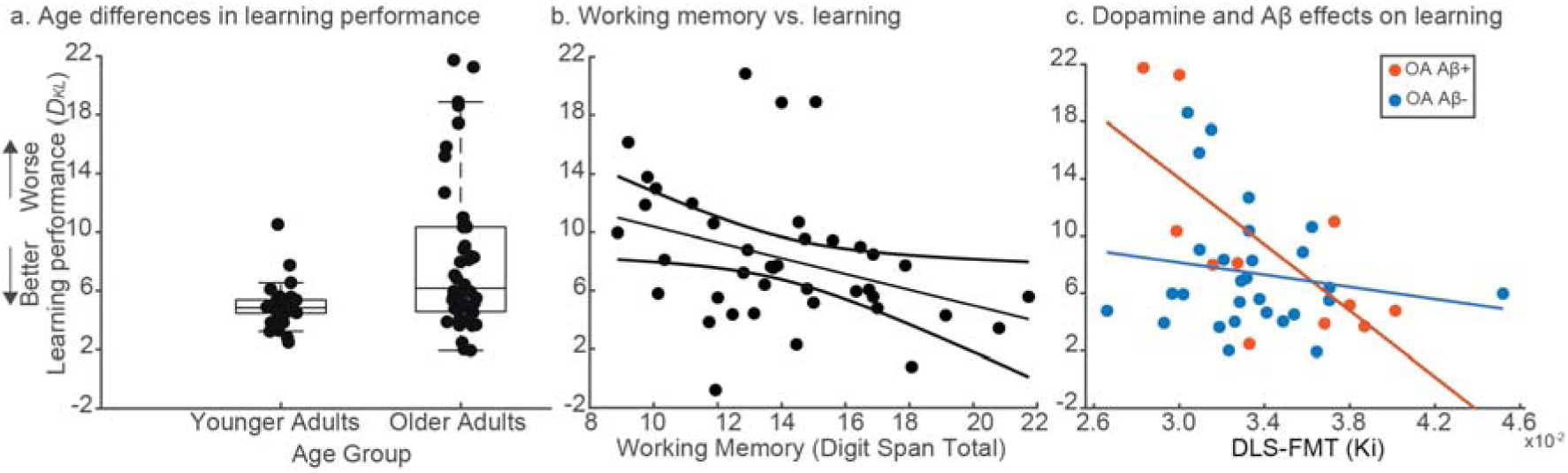
Learning Performance. a.) differences in learning performance between older and younger adults. b). relationship between learning performance and the digit span total score controlling for age in older adults. c.) relationship between learning performance and dorsolateral striatum (DLS) dopamine synthesis capacity for Aβ positive (red) and Aβ negative (blue) older adults.

To understand how the variability in older adult learning reflected broader cognitive abilities, we assessed its association with working memory and episodic memory in older adults. Age was included as a nuisance covariate within the older cohort. Learning performance was significantly associated with working memory (*Β* = −0.23, t = −2.3, p = 0.027), such that better working memory performance predicted more effective learning (**Figure 2b.)**. In contrast, episodic memory showed no significant relationship with learning (*Β* = 0.08, t = 0.61, p = 0.55).

We next modelled learning performance in older adults as a function of Aβ-PET status, DLS dopamine synthesis capacity, and temporal meta-region-of-interest tau-PET burden. Among the 33 participants with behavioural data and all three PET measures, multiple regression analysis revealed significant main effects of Aβ status (*Β*= −44.4, t = −2.5, p = 0.019) and DLS dopamine synthesis capacity (*Β*= −1446.9, t = −3.11, p = 0.0042), as well as a significant interaction between these variables (*Β*= 1241, t = 2.31, p = 0.028) such that dopamine showed a greater benefit for learning performance in Aβ positive than Aβ negative individuals. Temporal meta-ROI tau PET SUVR was not associated with learning performance (*Β*= −0.77, t = −0.13, p = 0.90). To maximise sample size, we re-estimated the model excluding the tau PET predictor, allowing inclusion of additional participants without tau PET (n = 39). This analysis showed a consistent pattern of results, with significant main effects of Aβ status (*Β*= −34.2, t = −2.32, p = 0.026), DLS dopamine synthesis capacity (*Β*= −1156.1, t = −3.31, p = 0.0022), and their interaction (*Β*= 943.7, t = 2.18, p = 0.036) (**Figure 2c.)**.

### Brain network activity and its relationship to learning, dopamine, and amyloid

We next investigated the cortical underpinnings of these behavioural findings. To assess this, we analysed the brain network responses to feedback throughout the learning task, probing how differential functional network activity to prediction errors (i.e. incorrect feedback) relates to learning, AD pathology, and DLS dopamine. We used spatial independent component analysis (ICA) to derive task-related activity within large-scale functional brain networks. Based on hypothesised network involvement, we extracted time series for each participant from the DMN and a frontostriatal network (**Supplementary Figure 2**). The ICA procedure yielded a canonical DMN with co-activations in the posterior cingulate, angular gyrus and prefrontal cortex (**Figure 3a., Supplementary Figure 2a.**). We also estimated a frontostriatal network capturing coactivations in the ventromedial prefrontal cortex and the ventral caudate (**Figure 3d., Supplementary Figure 2b.**). For each participant, we fitted a general linear model to the network time series and extracted parameter estimates contrasting trials where the participants response was incongruent (i.e. incorrect) versus congruent (i.e. correct) with the feedback received. Due to ceiling effects in learning performance and the poor coverage of FMT PET in younger adults, we restricted these analyses the older adults. In the full sample of 42 older adults, we observed that the difference of DMN activity to incorrect vs. correct feedback was significantly associated with individual learning performance (r(40) = −0.40, p = 0.009), indicating that greater DMN activity during incorrect feedback was associated with more effective learning (**Figure 3b**.). When investigating the effect of DLS dopamine synthesis capacity in the subset of 37 participants with FMT PET data, we observed a marginal association between differential frontostriatal feedback activity and dopamine synthesis capacity (r(35) = 0.33, p = 0.052), suggesting that greater endogenous DLS dopamine synthesis capacity tended to correspond with greater frontostriatal activity to incorrect feedback (**Figure 3e**.).

**Figure 3.**
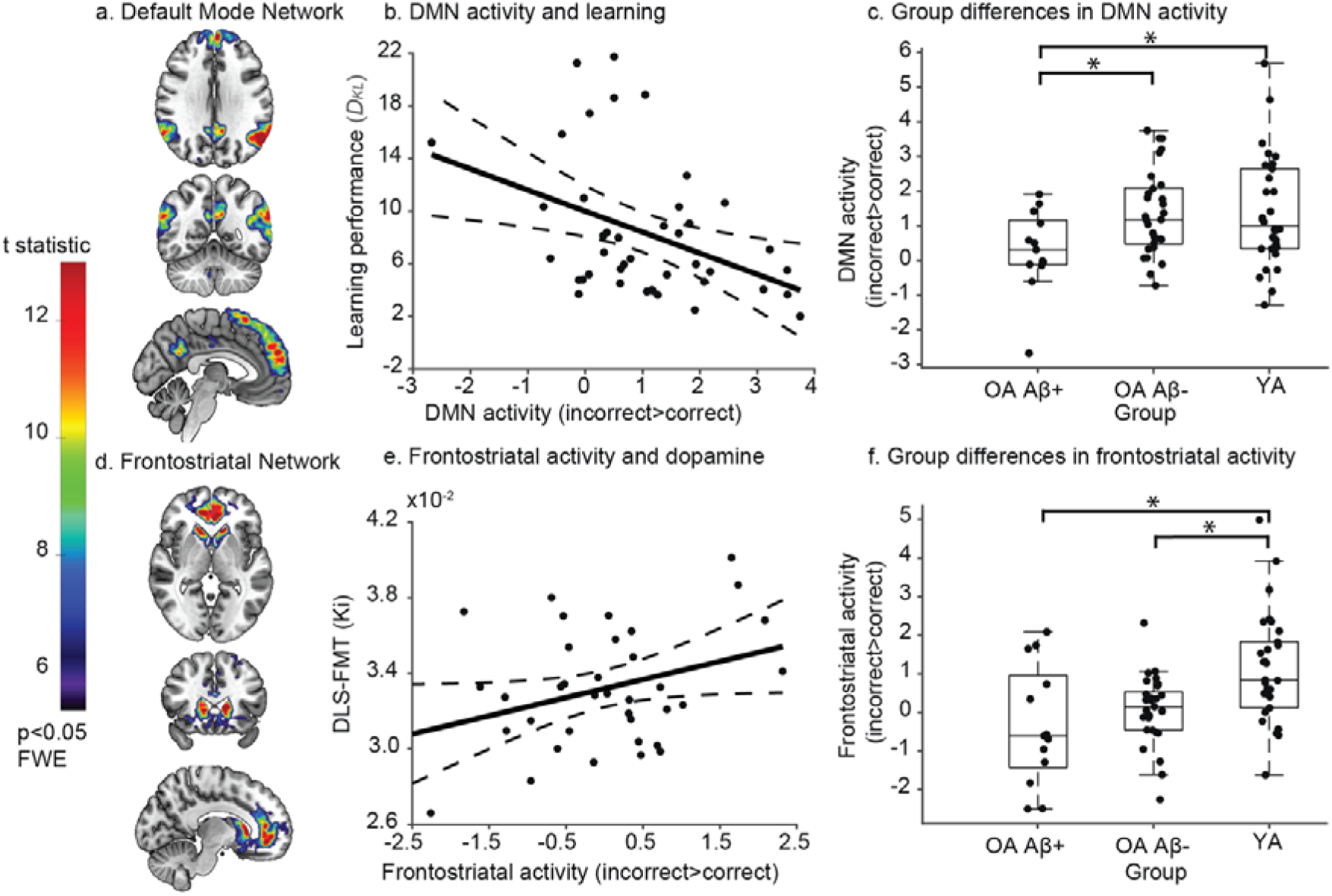
Brain network activity. a.) default mode network (DMN) spatial map estimated by spatial independent components analysis (ICA). b.) relationship between learning performance and DMN activation to incorrect minus correct trials in older adults (OA). c.) differences between DMN activation to incorrect minus correct trials for Aβ positive OA, Aβ negative OA, and young adults (YA). d.) frontostriatal network spatial map estimated by spatial ICA. e.) relationship between dorsolateral striatal (DLS) dopamine synthesis capacity and frontostriatal activation to incorrect minus correct trials in OA. f.) differences between frontostriatal activation to incorrect minus correct trials for Aβ positive OA, Aβ negative OA, and YA. * p<0.05

We next tested whether network responses differed as a function of Aβ status. Aβ-positive older adults showed attenuated modulation of DMN activity during incorrect feedback compared with Aβ-negative older adults (t(40) = −2.66, p = 0.011, Cohen’s d = −0.89) and younger adults (t(41) = −2.21, p = 0.033, Cohen’s d = −0.78). In contrast, modulation of DMN activity during incorrect feedback did not differ between Aβ-negative older adults and younger adults (t(57) = 0.047, p = 0.96, Cohen’s d = 0.012), suggesting that alterations in DMN activity to error related feedback signalling are driven by Aβ pathology rather than normative aging (**Figure 3c**.). In contrast, differential frontostriatal activity to incorrect feedback did not significantly differ between Aβ-positive and Aβ-negative older adults (t(40) = −1.08, p = 0.29, Cohen’s d = −0.32). However, a robust difference was present between Aβ-positive older adults and younger adults (t(41) = -2.98, p = 0.0048, Cohen’s d = -0.97), and Aβ-negative older adults and younger adults (t(41) = -3.36, p = 0.0014, Cohen’s d = -0.88) indicating strong age-related changes in frontostriatal error processing that were not specific to Aβ pathology (**Figure 3f**.).

### Effective connectivity to support learning through feedback modulation

We next used Dynamic Causal Modelling (DCM) to examine the system-level interactions underpinning the joint influence of Aβ and DLS dopamine on learning performance and their relationship to DMN and frontostriatal network connectivity. DCM allows average effective connectivity during the learning task (A matrix) and its stimulus-locked modulation during incorrect feedback (B matrix) to be inferred, thereby characterising how the functional system supporting learning responds to prediction error signals (i.e. incorrect feedback). These DCM analyses allow estimation of several key biophysical parameters of interest. First, the between network effective connectivity, which estimates whether one network increases (i.e. excitatory) or decreases (i.e. inhibitory) the activity of the other. Second, the self-inhibition parameters, which capture the intrinsic response or gain to extrinsic inputs. These parameters scale the default self-inhibition of a region (-0.5Hz) where a positive value indicates increased self-inhibition and a negative value indicates decreased self-inhibition (i.e. a relative shift toward excitation). Group-level inference on connectivity parameters and their association with Aβ and DLS dopamine was performed using Parametric Empirical Bayes (PEB) allowing us to infer how the different imaging markers influence the extrinsic or intrinsic processes that govern these brain network dynamics. In group-level PEB analyses interpretation was restricted to parameters with strong Bayesian evidence corresponding to posterior probabilities (Pp) greater than 0.99.

The DCM analysis comprised the DMN, frontostriatal network, and ventral visual network (**Supplementary Figure 3**), with the latter serving as the node receiving visual input during feedback trials (C matrix). A fully connected model with self-inhibitory connections was specified for the A matrix. For the B matrix, incorrect feedback trials were introduced as modulatory effects on connections between the DMN and frontostriatal network, as well as the self-connections of each region (**Figure 4a**.). This specification was motivated by our primary interest in the DMN’s response to incorrect feedback and its relationship to learning and Aβ. Frontostriatal connectivity was included to account for the parallel and interactive influence of DLS dopamine on learning. Connections between the visual cortex and frontostriatal network were excluded from the B matrix, as we had no a priori basis for expecting Aβ to influence learning via this pathway.

**Figure 4.**
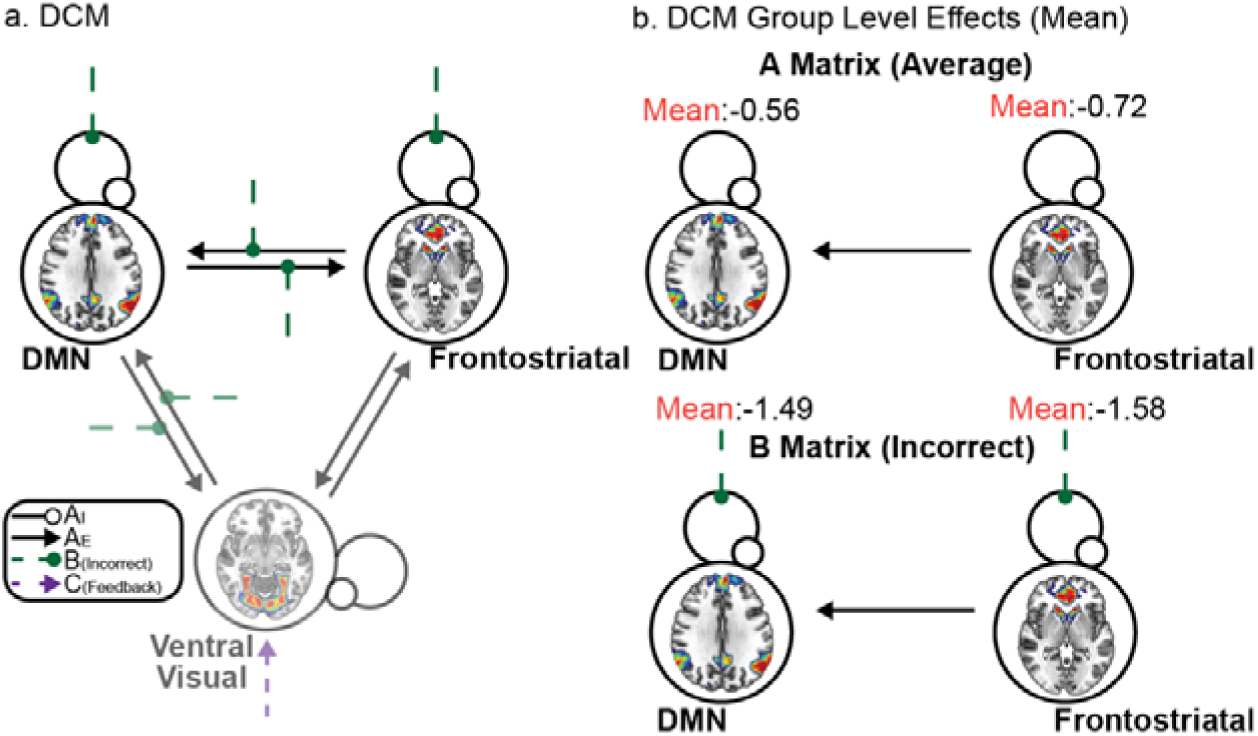
Dynamic Causal Model (DCM). a.) Fully connected DCM (black arrow) including self-inhibitory connections (black line with circle). The system receives inputs during feedback trials via the ventral visual network (purple arrow) with incorrect feedback providing a modulatory input to all connections to and from the default mode network (DMN) as well as the self-inhibitory connection of the DMN and frontostriatal networks (green line with circle). b.) Parametric empirical Bayes result indicating the mean (i.e. commonalities) effects. Top row shows the A matrix parameters representing the average effective connectivity during the task and the bottom row shows the B matrix representing the additive response to the A matrix during incorrect feedback. Self-connection parameters are the log of scaling parameters that multiply up or down the default self-connection (-0.5Hz), where positive values indicate greater self-inhibition, and negative values indicate less self-inhibition (i.e. a relative shift towards excitation). Red text indicates effects moving a connection towards excitation / dis-inhibition.

The DCM analysis for the 37 older adults with both PET and fMRI revealed small decreases in tonic self-inhibition within both DMN and frontostriatal networks throughout the task (DMN: *Β*= −0.56, Pp > 0.99; frontostriatal: *Β*= −0.72, Pp > 0.99), with feedback following incorrect trials substantially decreasing this same self-inhibition (DMN: *Β*= −1.49, Pp > 0.99; frontostriatal: *Β*= −1.58, Pp > 0.99). This indicates that error processing is characterised by a shift in DMN and frontostriatal gain resulting in a transient decrease in self-inhibition making them more responsive to extrinsic inputs during incorrect feedback (**Figure 4b**.).

Next, we investigated the role of Aβ and DLS dopamine synthesis capacity on the tonic effective connectivity between the DMN and frontostriatal network throughout the task (**Figure 5**). Aβ positivity was associated with inhibitory effective connectivity to the DMN from the frontostriatal network (*Β*= −0.34, Pp > 0.99). In contrast, higher DLS dopamine synthesis capacity was associated with an excitatory effect on this same connection (*Β*= 0.17, Pp > 0.99). This suggests opposing roles of DLS dopamine and Aβ on the tonic coupling between the DMN and frontostriatal network throughout the task (**Figure 5a**.).

**Figure 5.**
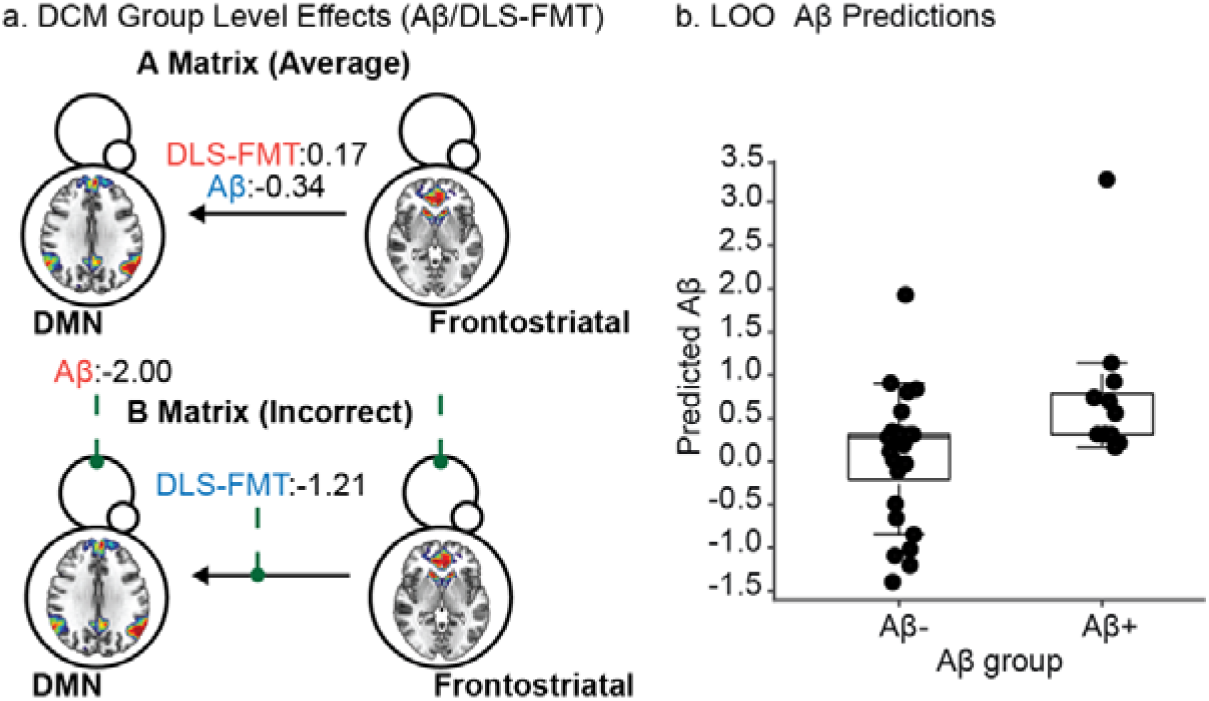
Influence of Aβ status and dorsolateral striatal dopamine on network interactions. a.) Parametric Empirical Bayes effects of Aβ status and dorsolateral striatal (DLS) dopamine synthesis capacity on each effective connectivity parameter. Top row shows the A matrix parameters representing effect of Aβ and (DLS) dopamine synthesis capacity on the average effective connectivity during the task. Bottom row shows the B matrix representing the effect of Aβ and (DLS) on to modulation of network effective connectivity during incorrect feedback. Red text indicates effects moving a connection towards excitation / dis-inhibition, and blue text indicates effects moving a connection towards inhibition / self-inhibition. Self-connection parameters are the log of scaling parameters that multiply up or down the default self-connection (-0.5Hz), where positive values indicate greater self-inhibition, and negative values indicate less self-inhibition (i.e. a shift towards excitation). b.) Leave one out (LOO) out of sample prediction of Aβ using the modulation of the DMN self-inhibition during incorrect feedback as the only predictor.

When investigating the impact of Aβ and DLS dopamine on the modulatory effect of incorrect feedback on the DMN and frontostriatal networks we observed further opposing influences on network dynamics. In response to incorrect feedback, we found that Aβ pathology induced a strong dis-inhibitory effect on the DMN (*Β*= −2.00, Pp > 0.99). This transient increase in dis-inhibition of the DMN during incorrect feedback (i.e. increases from normative dis-inhibition **Figure 4b**.) further shifts the DMN response to extrinsic inputs towards hyperexcitability, truncating the dynamic range of the DMN when processing prediction errors. Conversely DLS dopamine related to an increase in the inhibitory influence of the frontostriatal network to the DMN during incorrect feedback (*Β*= −1.21, Pp > 0.99). Suggesting that DLS dopamine exerts an opposing extrinsic inhibitory influence on the DMN when processing incorrect feedback (**Figure 5a**.).

Finally, we investigated how well the level of self-inhibition (observed as dis-inhibition **Figure 5a**) of the DMN during incorrect feedback predicted Aβ status using leave-one-out cross validation. In the full sample of older adults with Aβ-PET (n = 42) this single value of DMN self-inhibition (representing the excitation–inhibition (E/I) balance) during error processing to incorrect feedback reliably predicted Aβ out-of-sample (t(40)=2.542; p=0.015; Cohen’s d=0.851) (**Figure 5b**). This supports the finding that the E/I balance of the DMN during error processing is a sensitive functional marker of Aβ burden in cognitively normal older adults.

## Discussion

We show in cognitively normal older adults that feedback learning is impaired in response to Aβ burden. However, we found that higher levels of DLS dopamine synthesis capacity preserved normal learning performance in Aβ positive participants. Within these older adults we then model the cortical processing to prediction error signalling (i.e. feedback following an incorrect response) that supports this effective learning. We found that differential DMN activity to feedback reflects a participant’s learning capacity and that this modulation of DMN activity during prediction error processing is attenuated in Aβ positive participants. Through analysis of effective connectivity between the DMN and frontostriatal networks, we observed an Aβ related increase in the gain of the DMN during the processing of prediction errors and a compensatory influence of DLS dopamine on the coupling between the two networks. Our findings establish a systems-level framework where preclinical Aβ pathology impairs learning by disrupting the dynamic modulation to feedback of the DMN, and that this disruption can be mitigated by DLS dopamine function.

We show in cognitively normal older individuals that Aβ disrupts learning independently of temporal lobe tau, consistent with the notion that this learning is associated with executive functioning, rather than episodic memory. However, when investigating the role of striatal dopamine in older adults we find that increased DLS dopamine synthesis capacity may compensate for this Aβ related impairment to support effective feedback learning. This supports prior work implicating deficits in executive function as an indicator of Aβ related impairment in preclinical AD^11–13^, while providing support for the role of dopamine in promoting cognitive resilience to Aβ through improved feedback learning^18^. These findings complement parallel work examining the same paradigm and cohort, in which Morin et al.^26^ demonstrated that ventral striatal dopamine synthesis did not compensate for AD pathology to support episodic memory. When considered together these findings suggest that feedback learning and episodic memory, whilst engaging overlapping neural systems, are differentially vulnerable to AD pathological processes and draw on dopaminergic support in distinct ways. The resilience to Aβ afforded by elevated dopamine synthesis capacity may therefore be process and pathology specific, operating through learning via prediction error signalling during feedback^32^ rather than through mechanisms supporting the consolidation of episodic memories^33^, which are more dependent on medial temporal lobe function that is primarily compromised by tau^34^.

We show that Aβ attenuates the differential response of the DMN to feedback, and that this attenuation is associated with learning performance. This reduced DMN responsivity to afferent model updating signals is consistent with the well-documented reduction in deviant responses in mismatch negativity paradigms in Aβ-positive and AD participants^35–37^ likely reflecting cholinergic disruption^38^. Furthermore, our DCM analysis revealed that Aβ relates to self dis-inhibition of the DMN during incorrect feedback (i.e. hyperexcitability), reflecting a disruption in the dynamic responses of the DMN in Aβ-positive individuals when processing prediction errors. Prior work has demonstrated that high Aβ burden is associated with DMN hyperexcitability and failure to deactivate during stimulus repetition^39–43^ ; our findings extend this to the feedback learning domain. These results add to the literature linking Aβ to disrupted neuronal gain control and E/I balance^44–46^, providing behavioural evidence that these aberrant dynamics impair learning via model updating through prediction error signalling consistent with disrupted predictive coding mechanisms^4^.

Dopamine is known to modulate corticostriatal activity and inter-network connectivity with canonical brain networks, including the suppression of DMN regions^47–50^. Here, we show that DLS dopamine rebalanced the coupling between DMN and frontostriatal networks, opposing the effect of Aβ on the same connection. Furthermore, we found that DLS dopamine increased the inhibition of the DMN by the frontostriatal network during prediction errors, an opposing signal to the dis-inhibitory impact of Aβ on the DMN. The action of DLS dopamine to rebalance the effective coupling between the DMN and frontostriatal network in the face of Aβ may represent a critical compensatory mechanism to support learning via prediction errors consistent with prior work showing upregulated dopamine synthesis capacity and frontostriatal engagement in cognitively preserved older adults^19,51–53^.

Our findings suggest the DMN functions as part of a dynamic feedback-learning system supported by dopaminergic signalling through frontostriatal connectivity to restore learning capacity when burdened with Aβ pathology. This interpretation is consistent with emerging frameworks positioning the DMN at the intersection of value-based processing, prediction error signalling, and internal model updating via intrinsic function ^24,25^ as well as dynamic interactions with frontostriatal regions ^54–56^. When considered together, dopamine-dependent modulation of DMN during model updating may represent a key mechanism through which the brain maintains associative learning in the face of co-localised Aβ burden.

This work has several limitations. First, the sample size investigated here was relatively small, particularly when assessing the multivariate effects of Aβ, tau and FMT on learning. To address this, we restricted our analysis to interactions between the DMN, frontostriatal network, Aβ, and DLS dopamine and did not perform extensive exploratory brain wide analyses. In addition, we utilised Bayesian modelling approaches that ensure a parsimonious description of the data by weighting model evidence against complexity, reducing chances of overfitting. Notwithstanding, a sample of over forty cognitively normal older adults with task-based fMRI, Aβ, FMT, and tau PET in combination with in-depth neuropsychological assessment provides a rich sample to characterise the influences of Aβ and dopamine on brain functioning that underlies learning performance. Second, our analytical decisions and hypotheses were restricted to how prediction errors (i.e. incorrect feedback) modulate the effective connectivity within and between the frontostriatal network and DMN and how this relates to Aβ. This model omits the impact of tau and medial temporal activity which we have previously shown is related to predictive coding (i.e. repetition suppression) ^39^. However, our primary behavioural analysis did not show an effect of temporal meta-region of interest tau on learning, or a relationship between learning and episodic memory, implying that our learning metric of interest is not closely related to early tau accumulation or variability in cognition subserved by hippocampal function. Third, the task and behavioural learning index we derived does not capture all the nuances of probabilistic learning or reward-based value assignment. Furthermore, as the participants were instructed to identify the outcomes for a realtor (rather than the value on a given trial) the prediction error on a low probability response (i.e. losing on the winning realtor) should not shift the prior belief distribution of the associated realtor. As such, it is hard to unpack the specific pathological role of Aβ on model and precision updating. Finally, our approach to collapse learning across both loss and reward stimuli does not take into account the differential role of striatal dopamine in value assigned feedback. As our interest was in learning mechanisms linked to basic stimulus response associations we restricted our analysis of dopamine synthesis capacity to the DLS. The DLS has been implicated in stimulus response association^27,28^ and contrasts with the role of the dorsomedial striatum which has been implicated in more complex goal-oriented learning where probabilistic associations may vary over time as well as vary based on value assignment of feedback^57^. Additional investigations will be required to fully elucidate the complex role of striatal dopamine in supporting probabilistic feedback learning in preclinical AD.

## Conclusion

We show that Aβ impairs learning through functional changes in the DMN impeding the capacity of the brain to update in response to prediction errors. However, this detrimental action of Aβ on brain function is partially restored by the role of striatal dopamine acting on frontostriatal effective connectivity to the DMN. In this way dopamine may provide resilience to the effects of Aβ by rebalancing frontostriatal and DMN interactions to support feedback learning via effective model updating in response to prediction errors.

## Methods

### Participants

Fifty cognitively normal older adults and thirty healthy younger adults were recruited from the longitudinal Berkeley Aging Cohort Study. Participants were screened using a standard neuropsychological battery and included if performance fell within normative ranges (Mini-Mental State Examination ≥ 25 and all cognitive test scores within 1.5 standard deviations of normative means). All participants provided written informed consent in accordance with protocols approved by the Institutional Review Boards of the University of California, Berkeley and Lawrence Berkeley National Laboratory. All participants completed an fMRI scanning session during which they performed the probabilistic reward learning task (Real estate selling task). A subset of participants additionally underwent PET imaging to measure dopamine synthesis capacity using [^18^F]fluoro-L-m-tyrosine, tau burden using [^18^F]flortaucipir, and amyloid-β pathology using [^11^C]Pittsburgh Compound-B.

### Real estate selling task design

During scanning, participants completed the Real Estate Selling Task (**Figure 1**), a probabilistic learning paradigm designed to examine feedback-based learning. This task and sample have been previously described (see Morin et al.^26^ ). Here, we only interrogate learning from the first two runs of the task and do not analyse the learning data for the final two runs that have new stimulus outcome associations. Participants were instructed that they owned a portfolio of houses and were collaborating with three real estate agents to sell them. The agents differed in their typical performance with one usually selling houses for a profit (“reward”), one consistently breaking even (“neutral”), and one usually selling houses at a loss (“loss”). Participants were instructed to learn the identity of each realtor based on feedback. Although participants received monetary feedback on each trial, trial outcomes were predetermined by the experimenter and were not contingent on participant responses.

### Trial structure

Each trial began with a cue (2.0 s) showing an image of one realtor. Participants pressed one of three buttons to indicate whether they believed the realtor corresponded to the reward, neutral, or loss agent. This cue period was followed by a jittered delay (2.4–4.8 s). Next, the target phase (3.0 s) presented an image of the house being sold. The house image was outlined in a colour indicating the sale outcome: green (profit), grey (break-even), or red (loss). A feedback screen (1.0 s) then displayed the monetary outcome (+$1 for reward trials, –$0.50 for loss trials, or “=” for neutral trials). Trials were separated by a jittered inter-trial interval (2–10 s). If participants failed to respond during the cue period, the target image appeared with a question mark overlay and no outcome was shown. These trials were omitted from analyses.

### Outcome contingencies

Outcome probabilities were fixed for the three realtors, two were probabilistic (reward and loss) and one was deterministic (break-even). The reward realtor produced profits on 80% of trials and losses on 20%. The loss realtor produced losses on 80% of trials and profits on 20%. The neutral realtor produced break-even outcomes on 100% of trials. Participants were not informed of these probabilities and were instructed to categorize each realtor based on typical performance rather than predict the outcome of a particular sale. Participants completed two runs of the task (42 trials per run; 84 trials total).

### Learning Index

To quantify participant-specific learning of the probabilistic stimulus–outcome contingencies, we computed a trial-wise learning index using Kullback–Leibler Divergence (KL divergence). The KL divergence measures the difference between the distribution of outcomes previously experienced for a given realtor and the participants response distribution to the same realtor. This measure provides a scalar value per trial for how well each participants choices followed the true outcome statistics for each stimulus.

For each participant, behavioural data were aligned so that the presentation and response categories for feedback were classified into three categories corresponding to reward, neutral, and loss feedback. For each trial *t*, outcome distributions were computed separately for each stimulus identity (i.e., realtor). To define the outcome distribution *P*, the distribution of outcomes previously observed for that stimulus up to trial *t* − 1 was calculated. To define the participant response distribution *R*, the distribution of participant responses up to trial *t* was calculated. Both distributions were normalised to probability distributions and smoothed using Laplace regularisation (ε = 0.1) to avoid undefined values when probabilities approached zero. The trial wise KL divergence was calculated as the divergence between these two distributions

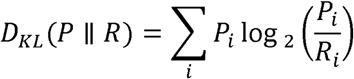

This value reflects the mismatch between the participant’s response model for a realtor and the observed outcomes for the realtor.

Trial-wise KL divergence values were computed separately for each stimulus category (reward, neutral, loss). To capture cumulative learning across the task, the trial wise KL divergence for each realtor was integrated across trials using trapezoidal integration in MATLAB. Because reward and loss stimuli had probabilistic outcomes whereas neutral stimuli were deterministic, we averaged the integral of KL divergence for probabilistic conditions (reward and loss stimuli) and used this as our scalar index of participant learning. These learning indices were calculated across the task (runs 1 and 2), resulting in participant-level measures reflecting the cumulative divergence between outcomes and participant responses. Lower KL divergence values indicate that the participant’s response distribution more closely matched the outcome distribution, reflecting more accurate learning of stimulus–outcome contingencies, whereas higher values indicate greater divergence between response and outcome contingencies (see **Supplementary Figure 1**).

### PET Imaging

#### Dopamine synthesis capacity

Dopamine synthesis capacity was measured using the irreversible PET tracer [^18^F]fluoro-L-m-tyrosine. Approximately 2.5 mCi of tracer was injected intravenously, and participants were scanned for 90 minutes on a Siemens Biograph TruePoint 6 PET/CT scanner. Participants received oral carbidopa (2.5 mg/kg) one hour before scanning to reduce peripheral metabolism.

Dynamic frames were reconstructed using ordered subset expectation maximization with attenuation and scatter correction and smoothed with a 4 mm FWHM kernel. Ki maps representing dopamine synthesis capacity were computed using Patlak graphical analysis with posterior cerebellar grey matter as the reference region. As our learning effects of interest are related to how well a participant learns stimulus response associations (i.e. if a realtor is reward/loss/break even) we focussed on the PET signal from the dorsal putamen to capture dorsolateral striatum dopamine synthesis capacity. Striatal regions of interest (ROIs) were manually drawn on each participant’s T1-weighted structural scan as previously described^58^ using Mango software.

### Amyloid and tau imaging

Amyloid-β burden was measured in a subset of participants using [^11^C]Pittsburgh Compound-B PET, and tau burden was measured using [^18^F]flortaucipir PET. PiB distribution volume ratios were computed using Logan graphical analysis with cerebellar grey matter as the reference region. Tau PET standardised uptake value ratios were calculated from tracer uptake between 80–100 minutes post-injection normalized to inferior cerebellar grey matter.

### MRI Data Acquisition

MRI data were acquired on a 3T Siemens TIM/Trio scanner located at the Henry H. Wheeler Jr. Brain Imaging Center using a 32-channel head coil. High-resolution T1-weighted structural images were acquired using a magnetization-prepared rapid gradient echo (MPRAGE) sequence (1 mm isotropic voxels; TR = 2300 ms; TE = 2.98 ms; matrix = 256 × 240; 160 sagittal slices). Functional images were collected using a multiband T2*-weighted echo-planar imaging sequence (slice acceleration factor = 4; TR = 2400 ms; TE = 36 ms; flip angle = 45°). Eighty-eight slices covering the whole brain were acquired with 1.54 mm isotropic voxels (matrix = 138 × 138). A total of 246 volumes were acquired per run and two runs were collected for each participant. Opposite phase-encoded EPI fieldmaps (anterior-posterior and posterior-anterior) were collected for distortion correction.

### fMRI Preprocessing

MRI preprocessing was performed using fMRIPrep (version 22.0.2) ^31^, implemented through Nipype. Structural images were corrected for intensity non-uniformity using N4BiasFieldCorrection, skull stripped using the ANTs brain extraction workflow with the OASIS30ANTs template, and segmented into grey matter, white matter, and CSF using FSL FAST. Cortical surfaces were reconstructed using FreeSurfer. Functional preprocessing included slice-time correction (AFNI 3dTshift), head-motion estimation using MCFLIRT, susceptibility distortion correction using phase-encoded fieldmaps, and co-registration of functional images to the T1 reference using boundary-based registration. Data were normalized to the MNI152NLin2009cAsym template. Resampling was performed using ANTs with Lanczos interpolation to minimize smoothing. The first three volumes of each run were discarded, and additional pre-processing steps including linear detrending, voxel wise grand mean scaling, and spatial smoothing (3 mm FWHM Gaussian kernel) were performed in MATLAB.

### Independent Component Analysis

Group spatial independent component analysis (ICA) was performed using the Group ICA fMRI Toolbox (GIFT, https://trendscenter.org/software/gift/). Pre-processed data from both younger and older adults were entered into the analysis to obtain robust estimates of task-related cortical networks. The imaging data for each run was treated as a separate session for each subject. The ICA dimensionality was set to 50 components. A two-stage dimensionality reduction procedure using principal component analysis was performed first at the subject level and then at the group level. Components were estimated using the Infomax algorithm and the ICA procedure was repeated 20 times with component stability evaluated using ICASSO. Subject-specific spatial maps and time courses were obtained through back-reconstruction. Component time courses were normalized within each participant and run (z-scored) to allow comparisons across subjects. Group-level spatial maps were obtained using one-sample t-tests on the back-reconstructed spatial maps to identify voxels significantly contributing to each component (p<0.05 FWE). Three components were selected a priori for further analyses, a canonical default mode network (DMN) and a frontostriatal network which were hypothesised to be involved in stimulus–response learning, and a ventral visual component hypothesised to be responsive to the visual stimuli.

### ICA-Based GLM Analysis

Task-related responses within the DMN and frontostriatal networks were modelled using a general linear model applied to the normalised ICA time series. The feedback phase of the task was modelled with stick functions convolved with the hemodynamic response function to form regressors corresponding to correct and incorrect trials (defined as participant responses congruent or incongruent with the feedback outcome). Parameter estimates (β-weights) for each condition were extracted and contrasted (correct minus incorrect feedback) to characterise network responses to feedback during learning. Regressors of no interest included spike regressors for motion outliers (framewise displacement > 0.5 mm), the six motion parameters and the derivatives of the motion parameters. The GLM was fit on each run independently and parameter estimates averaged across the runs. For subsequent system-level analyses, nuisance effects were removed from the component time courses via linear regression and the residual time series for each run concatenated together.

### Dynamic Causal Modelling

Effective connectivity between networks during task performance was examined using deterministic bilinear dynamic causal modelling implemented in SPM12^59^. Three nodes were included in the model, the DMN, the frontostriatal network, and a ventral visual network that served as an input node receiving sensory information during feedback presentation.

DCM models neuronal population dynamics using a bilinear differential equation:

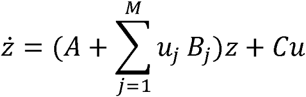

where *z* represents the neuronal state vector, *A* defines intrinsic coupling between regions, *B_j_* encodes modulatory effects of experimental inputs, and *C* specifies direct driving inputs.

Subject-level models were constructed following procedures described by Zeidman et.al^59^. A template DCM was created in the SPM graphical interface and subsequently populated with subject-specific ICA time series and task design matrices. The network architecture consisted of a fully connected bidirectional graph including self-connections for each node (A matrix). Feedback events served as driving inputs to the ventral visual node (C matrix). Based on the observed association between DMN activity and learning performance, incorrect feedback trials were included as modulatory inputs affecting connections to and from the DMN as well as the DMN self-connection (B matrix). Incorrect trials were coded as 1 and correct trials as 0 and subsequently mean-centred. This transformation ensured that the intrinsic connectivity parameters (A matrix) represented mean connectivity across the task, while modulatory parameters (B matrix) represented deviations from this baseline in response to incorrect feedback.

Within the DCM framework, connection parameters correspond to rate constants governing the change in neuronal activity in response to afferent signals. Positive values indicate excitatory influences (reduced damping), whereas negative values indicate inhibitory influences (stronger damping). Self-connections represent log-scaling parameters controlling local gain (i.e. E/I balance) that multiply up or down the default self-connection (-0.5Hz). Positive values indicate greater self-inhibition, and negative values indicate greater dis-inhibition (i.e. a relative shift towards excitation).

### Parametric Empirical Bayes

Group-level effects on effective connectivity were estimated using the Parametric Empirical Bayes (PEB) framework implemented in SPM. PEB uses hierarchical Bayesian modelling to estimate group-level effects while accounting for between-subject variability in connection strengths^60^. The second-level design matrix included (1) a constant term, (2) z-scored dorsolateral striatal dopamine synthesis capacity measured using [^18^F]fluoro-L-m-tyrosine PET, and (3) mean-centred Aβ status measured with [^11^C]Pittsburgh Compound-B PET.

Bayesian model reduction was applied to iteratively prune parameters that did not contribute to model evidence (Free Energy). Model evidence reflects a trade-off between accuracy (model fit to observed BOLD data) and complexity (number and magnitude of parameters). Posterior probabilities were computed by comparing models in which parameters were present versus absent. Parameters with posterior probability greater than 0.99 were considered to have strong Bayesian evidence. Primary analyses focused on modulatory parameters affecting coupling between the DMN and frontostriatal networks and the DMN self-connection during incorrect feedback. A secondary PEB model excluding dopamine synthesis capacity was estimated to maximize sample size for analyses of Aβ effects.

### Cross-Validation

To evaluate whether effective connectivity parameters predicted Aβ status, leave-one-out cross-validation was performed within the PEB framework. The modulatory parameter governing the DMN self-connection was used to predict out-of-sample Aβ status for each participant. Predicted and observed group labels were compared using a two-sample t-test.

## Acknowledgments

Avid Radiopharmaceuticals enabled the use of the 18F-Flortaucipir tracer but did not provide direct funding and were not involved in data analysis or interpretation. J.G. is supported by the Alzheimer’s Association (23AARF-1026883). W.J. is supported by the NIH (AG034570 and AG062542). M.B. is supported by NHMRC (APP1152623 and APP2008612).

## Author contributions

Conceptualization, J.G., W.J.J., and M.B.; formal analysis, J.G., T.M., H.C., and A.S.B.; data curation, J.G., T.M., H.C., A.S.B., and W.J.J.; methodology, J.G. and M.B.; writing – original draft, J.G., W.J.J., and M.B.; writing – reviewing & editing, J.G., T.M., H.C., A.S.B., W.J.J., and M.B.; supervision W.J.J. and M.B.

## Inclusion and diversity

We support inclusive, diverse, and equitable conduct of research.

