## Supplementary for "Dopamine Compensates for Amyloid-Induced Default Mode Network Dysfunction to Support Learning"

**Supplementary Tables**

|  | **Young Adults** | **OA With Aβ** | **OA With Aβ & FMT** | **OA With Aβ & fMRI** | **OA With Aβ & fMRI & FMT** | **OA With Aβ & FMT & FTP** |
| --- | --- | --- | --- | --- | --- | --- |
| **Age [Mean ± SD]** | [25.0, 3.72] | [76.07, 5.73] | [76.03, 5.82] | [76.24, 5.57] | [76.22, 5.65] | [75.67, 6.15] |
| **Education [Mean ± SD]** | [16.24, 1.79] | [16.89, 1.78] | [16.79, 1.73] | [16.88, 1.81] | [16.78, 1.77] | [16.76, 1.84] |
| **Sex [F / M]** | 12 / 18 | 25 / 19 | 22 / 17 | 23 / 19 | 20 / 17 | 18 / 15 |
| **MMSE [Mean ± SD]** | N/A | [28.71, 0.99] | [28.74, 1.02] | [28.70, 0.99] | [28.73, 1.02] | [28.76, 1.00] |
| **Aβ Status [+ / −]** | N/A | 13 / 31 | 11 / 28 | 13 / 29 | 11 / 26 | 7 / 26 |
| **Temporal Meta-ROI**  **Tau SUVR [Mean ± SD]** | N/A | N/A | N/A | N/A | N/A | [1.30, 0.15] |

**Supplementary Table 1 Sample characteristics.** OA = Older Adults; FMT = fluoromethyltyrosine PET; fMRI = functional MRI; FTP = flortaucipir PET; Aβ = amyloid-beta; MMSE = Mini-Mental State Examination; SUVR = standardised uptake value ratio. Values in brackets denote [Mean, SD].

**Supplementary Figures**

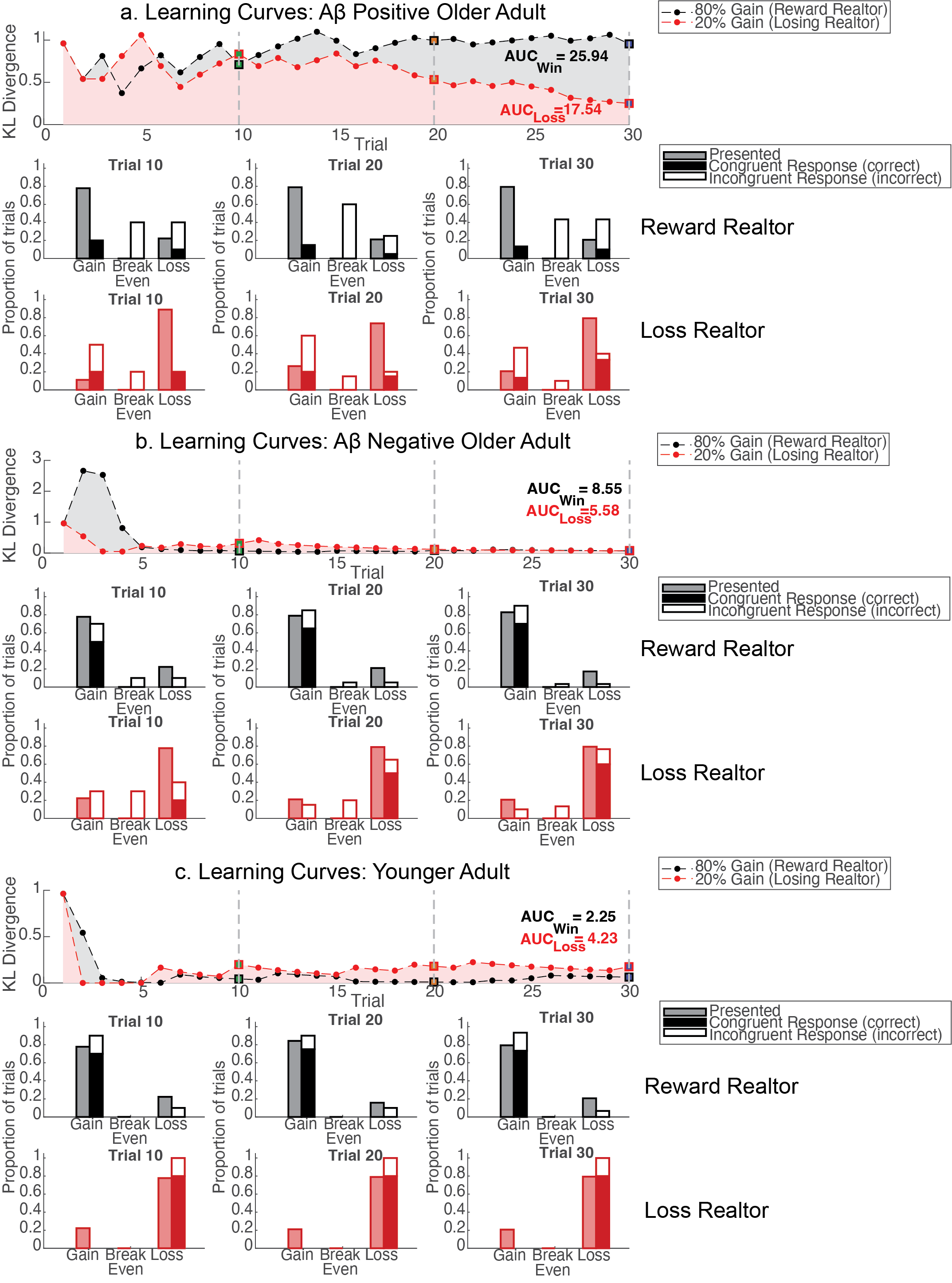

**Supplementary Figure 1 Representative learning curves and response vs. presented distributions throughout learning**. Top panels show KL divergence between the presented feedback distribution and the subject's response distribution over trials for an Aβ positive older adult (**a.**), an Aβ negative older adult (**b.**), and a younger adult (**c.**). Lower KL divergence indicates greater alignment between the subject's response distribution and the presented outcomes, reflecting improved learned associations. Shaded regions represent the integral of the learning curve, where a smaller value indicates faster and more complete learning across trials with the average integral for the reward and loss relator used as each subject’s metric for learning. The black and red curves show how the learned association between gaining or losing money evolve for the reward and loss realtor respectively. To illustrate how this metric relates to the concordance between responses and feedback, we sampled response and presentation distributions and at checkpoint trials (10, 20, 30). Lower panels display grouped bar charts at each checkpoint for the faces of the reward relator (80% gain) (middle row) and the loss relator (20% gain) (bottom row). Left shaded bars show the proportion of each feedback outcome (gain, break even, loss) presented up to that checkpoint; right bars show the subject's response distribution as a stacked bar, with the filled (condition-coloured) segment reflecting responses congruent (i.e. correct) with the received feedback on that trial, and the white segment reflecting incongruent (i.e. incorrect) responses.

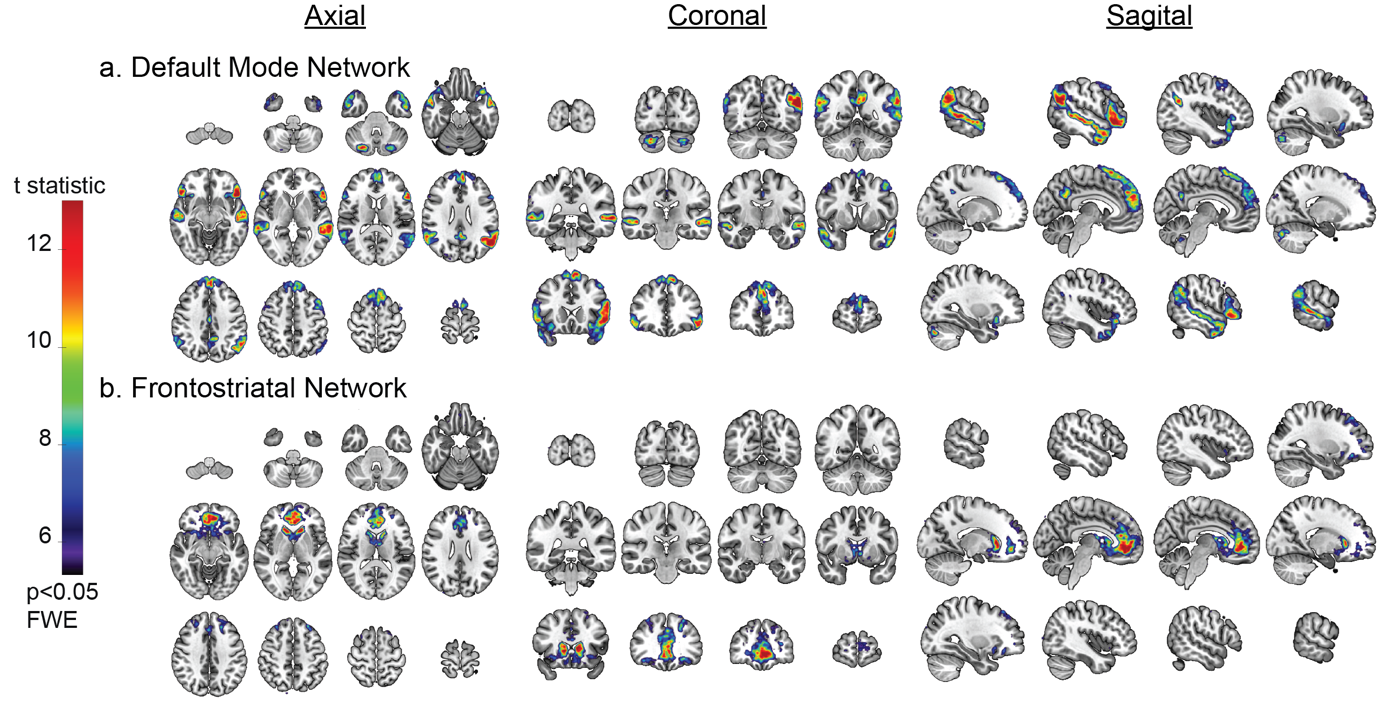

**Supplementary Figure 2** **ICA component maps**. Group t-maps of back reconstructed ICA spatial maps projected onto the MNI brain. Colour scale indicates the t-statistic of the group test against zero for the a.) Default mode network and b.) Frontostriatal network. Maps are thresholded at p<0.05 FWE corrected.

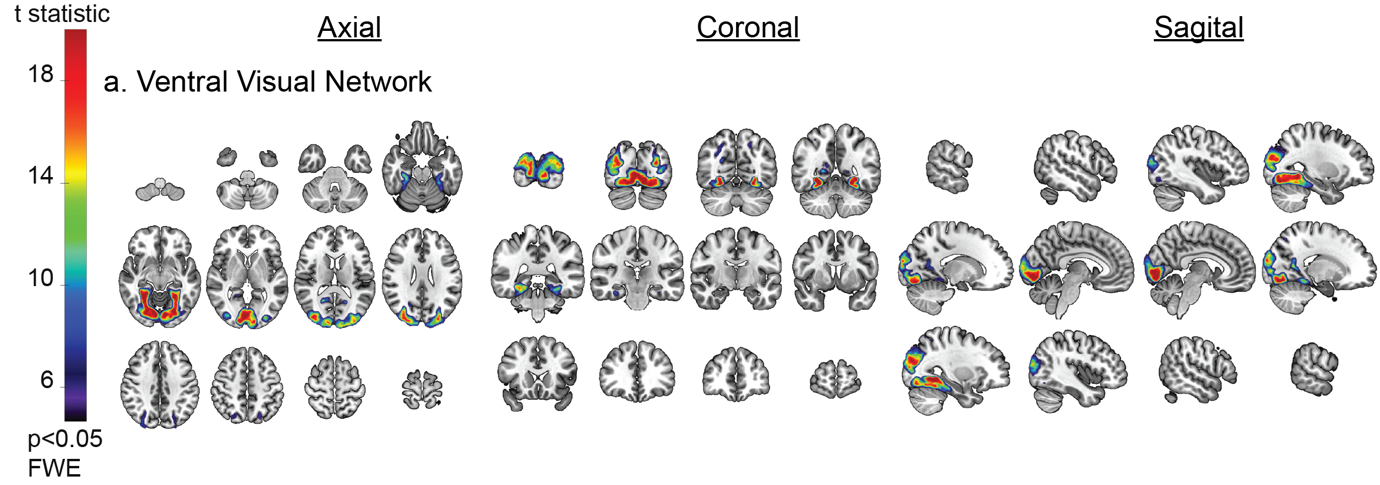

**Supplementary Figure 3 ICA component maps**. Group t-maps of back reconstructed ICA spatial maps projected onto the MNI brain. Colour scale indicates the t-statistic of the group test against zero for the a.) Ventral visual network. Maps are thresholded at p<0.05 FWE corrected.

**Supplementary Methods**

**Task Instructions:**

In this game, you are in the business of selling houses. There are 3 real estate agents that you are working with, but they are not equally skilled. One agent is more likely to lose you money when selling houses. One agent always breaks even (you neither gain money nor lose money). One agent is more likely to make you money when selling houses. Your goal is to figure out which agent is which.

First, you will see a picture of the agent. You must indicate whether you think they will make you lose money (‘b’), break even (‘n’) or gain money (‘m’) in this deal. At first, you will just have to guess.

Next, you will see a picture of the house they are selling for you along with feedback on how the sale went.

If the agent lost you money, there will be a red outline around the house and a display showing -$0.50. The unskilled agent will usually lose you money (approximately 80% of the time), so their houses will usually have the red box around them.

If the agent broke even (meaning you neither gained nor lost money), there will be a gray outline around the house and a display showing an equal sign.

If the agent made you money, there will be a green outline around house and a display showing $1. The skilled agent will usually make you money (approximately 80% of the time), so their houses will usually have the green box around them.

From this feedback, you will learn which agent is which.

Remember: The agent is what determines whether you gain or lose money, not the house.

It is important that when you see the real estate agent, you indicate whether you think they are the unskilled agent that tends to lose you money, the break even agent, or the skilled agent that tends to gain you money. Please respond as quickly as possible without sacrificing accuracy.

The skilled and unskilled agents don’t make you make/lose money 100% of the times. Sometimes the unskilled agent gets lucky and sometimes the skilled agent has a bad day. You should not be discouraged if your prediction about the money outcome with them is wrong, in a specific deal. Do your best to figure out which agent is which.

We know this is a just a laboratory experiment which is not like the real world. In order to make it a little more realistic, we have made the monetary outcomes real. The unskilled agent is making you lose real money, and the skilled agent is making you win real money. However, don’t worry: In our experiment, it is not possible for you to lose money overall. In other words, we will not take money away from you. The outcome is dependent on the realtor's skill level, not your guess. Your goal, during the experiment, is to correctly guess which outcome each realtor is usually associated with.
